# Impact of acquisition and modeling parameters on test-retest reproducibility of edited GABA+

**DOI:** 10.1101/2023.01.20.524952

**Authors:** Kathleen E. Hupfeld, Helge J. Zöllner, Steve C. N. Hui, Yulu Song, Saipavitra Murali-Manohar, Vivek Yedavalli, Georg Oeltzschner, James J. Prisciandaro, Richard A. E. Edden

## Abstract

Literature values for within-subject test-retest reproducibility of gamma-aminobutyric acid (GABA), measured with edited magnetic resonance spectroscopy (MRS), vary widely. Reasons for this variation remain unclear. Here we tested whether sequence complexity (two-experiment MEGA-PRESS versus four-experiment HERMES), editing pulse duration (14 versus 20 ms), scanner frequency drift (interleaved water referencing (IWR) turned ON versus OFF), and linear combination modeling variations (three different co-edited macromolecule models and 0.55 versus 0.4 ppm spline baseline knot spacing) affected the within-subject coefficient of variation of GABA + macromolecules (GABA+). We collected edited MRS data from the dorsal anterior cingulate cortex from 20 participants (30.8 ± 9.5 years; 10 males). Test and retest scans were separated by removing the participant from the scanner for 5-10 minutes. Each acquisition consisted of two MEGA-PRESS and two HERMES sequences with editing pulse durations of 14 and 20 ms (referred to here as: MEGA-14, MEGA-20, HERMES-14, and HERMES-20; all TE = 80 ms, 224 averages). Reproducibility did not consistently differ for MEGA-PRESS compared with HERMES or for 14 compared with 20 ms editing pulses. A composite model of the 0.9 and 3 ppm macromolecules (particularly for HERMES) and sparser (0.55 compared with 0.4 ppm) spline baseline knot spacing yielded generally better test-retest reproducibility for GABA+. Replicating our prior results, linear combination modeling in Osprey compared with simple peak fitting in Gannet resulted in substantially better test-retest reproducibility. These results highlight the importance of model selection for edited MRS studies of GABA+, particularly for clinical studies which focus on individual patient differences in GABA+ or changes following an intervention.

## 1. Introduction

Gamma-aminobutyric acid (GABA), the brain’s primary inhibitory neurotransmitter, is an important metabolite for normal brain function. GABA levels can be quantified *in vivo* noninvasively using proton magnetic resonance spectroscopy (^1^H MRS). However, due to its low concentration within the brain and overlapping spectral signatures of higher concentration metabolites, spectral editing is required for accurate *in vivo* quantification of GABA at 3 Tesla. Prior work has reported differences in brain GABA levels across the healthy lifespan^1–3^, as well as in various psychiatric^4,5^ and neurodegenerative conditions^6–8^.

One common spectral editing technique for GABA is MEscher-GArwood Point RESolved Spectroscopy (MEGA-PRESS^9^), which requires a relatively lengthy (e.g., 10-minute) acquisition to target one metabolite (e.g., GABA). Newer sequences allow for simultaneous editing of multiple low-concentration metabolites; for instance, the Hadamard Encoding and Reconstruction of MEGA-edited Spectroscopy (HERMES) sequence permits simultaneous detection of both GABA and glutathione (GSH, one of the most abundant antioxidant sources in the central nervous system) using a Hadamard-encoded editing scheme^10^. Due to the finite selectivity of the editing pulses, the edited 3-ppm GABA signal is confounded by a co-edited macromolecular (MM) resonance. Since these GABA and MM signals cannot be reliably separated, their composite ‘GABA+’ is commonly reported.

Reports of intrasubject variability in MRS-measured GABA+ vary widely. Reported test-retest within-subject coefficients of variation (CVs) vary from 1% to almost 30%^3,11–15^. Specific reasons for this wide range of reproducibility remain unclear. Maximizing test-retest reproducibility is becoming increasingly critical, for example as edited GABA+ measures are being used as an outcome measure for clinical trials^16,17^.

Several acquisition factors, including editing pulse duration and editing scheme complexity (MEGA-PRESS versus HERMES) may contribute to intrasubject variability in GABA+ measurements. Longer editing pulses are more selective (narrower bandwidth) and thus result in smaller contributions from co-edited MM signals^18^ and greater sensitization to motion and scanner frequency drift^19,20^. The extent to which 14 versus 20 ms editing pulses for MEGA-PRESS and HERMES sequences affect intrasubject reproducibility of GABA+ measurements has not been explicitly reported. HERMES interleaves four sub-experiments into the same acquisition to simultaneously edit for GABA+ and GSH^10^ compared to the two-step MEGA-PRESS scheme. One key part of post-processing is the alignment of sub-spectra before calculating Hadamard combinations in order to reduce subtraction artifacts in the GABA- and GSH-edited difference spectra^21–23^. Aligning four HERMES sub-spectra which lack a common strong ‘reporter signal’ might be more challenging and variable than aligning two MEGA sub-spectra that share the same residual water peak. Though we previously reported an intrasubject CV of 9.9% for GSH measured with HERMES, compared with 8.8% for GSH measured with MEGA-PRESS^14^, the extent to which HERMES might increase intrasubject variability of GABA+ is unclear.

Another potential factor influencing reproducibility is scanner frequency drift. Edited sequences such as MEGA-PRESS and HERMES are sensitive to scanner frequency drift^20,24^, caused by instability of the superconducting magnet, gradient heating or cooling, or subject motion that can differ between individual scanners^20,25^. This B_0_ instability may lead to increased linewidths and imperfect subtraction of signals in the OFF and the ON spectra in edited MRS experiments, but also changes to editing efficiency. In order to address B_0_ frequency drift during acquisition, a prospective frequency correction method called interleaved water referencing (IWR) was proposed and demonstrated to reduce between-subject variance in MM-suppressed GABA measurements^24^. Here we investigate whether IWR impacts intrasubject variability of GABA+.

Though our initial goals for this work were to characterize the effect of acquisition factors on GABA+ intrasubject variability, we decided *post hoc* to also examine the effects of modeling choices on GABA+ reproducibility. Our recent work^14^ reanalyzing the test-retest dataset reported in Prisciandaro et al. (2020)^12^ found that linear-combination modeling with the Osprey software package^26^ significantly improved HERMES GSH intrasubject reproducibility over simple peak fitting in Gannet^27^, reducing the within-subject CV from 19 to 9.9%. Here, in addition to replicating our finding of the benefit of Osprey over Gannet processing^14^, we aimed to examine how Osprey linear-combination modeling decisions affect GABA+ reproducibility. Among 61 GABA-edited MEGA-PRESS datasets with TE = 68 ms, we previously found that sparser spline knot spacing and inclusion of co-edited MMs in the linear combination model basis function significantly improved intersubject CVs^28^. Although the spline baseline is needed to account for lipid contamination and poor water suppression, it can also be a potential source of overfitting if given too many degrees of freedom. Highly flexible baseline models are likely to absorb substantial portions of the edited 3-ppm GABA+ signal, leading to overall lower and more variable estimates. Sparser spline knot spacing (0.55 ppm over 0.4 and 0.25 ppm) better estimated the background in the GABA-edited difference spectra for TE = 68 ms edited GABA+ data; here, we investigate if the same is true for TE = 80 ms GABA+ data. Linear combination modeling of GABA-edited spectra is improved by modeling of the co-edited MM signal at 3 ppm (MM_3co_)^28^. Here we fit three different models that yielded the lowest intersubject CVs in the recommended wide (0.5-4 ppm) modeling range with sparse (0.55 ppm) knot spacing in our previous work^28^, referred to in Osprey as ‘1to1GABA’, ‘1to1GABAsoft’, and ‘3to2MM’ (described in detail in the Methods). In the present work, we examined how these modeling decisions affected intrasubject reproducibility of GABA+ at TE = 80 ms, which has lower SNR due to reduced editing efficiency and greater T_2_ relaxation of GABA and MM signals compared with TE = 68 ms GABA+ data and subtly different signal lineshapes. For completeness, we also examined the impact of acquisition and analysis factors on intrasubject reproducibility of HERMES-measured GSH estimates. We aimed to replicate our finding of better GSH test-retest reproducibility using Osprey compared with Gannet processing^14^ and to determine whether editing pulse duration or dense versus sparse spline baseline knot spacing differentially affects GSH compared with GABA+ reproducibility.

In summary, in the present work acquired a new test-retest dataset to investigate the effects of sequence complexity (MEGA-PRESS versus HERMES), editing pulse duration (14 versus 20 ms), scanner frequency drift (IWR-ON versus IWR-OFF), spline baseline knot spacing (0.40 versus 0.55 ppm), MM model (1to1GABA, 1to1GABAsoft, and 3to2MM), and use of linear-combination modeling (Osprey versus Gannet simple peak fitting) on the test-retest reproducibility (within-subject CV) of GABA+. We predicted that shorter editing pulse duration (14 ms) and less sequence complexity (MEGA-PRESS) would result in better GABA+ test-retest reproducibility (lower within-subject CVs). We examined the effects of Osprey modeling choices *post hoc* and thus did not have hypotheses for these exploratory aims; however, in line with our previous work^14^, we predicted that Gannet processing would result in poorer reproducibility compared with Osprey.

## 2. Methods

### 2.1 MRS Acquisition

Twenty healthy adults (mean age: 30.8 ± 9.5 years; 10 males) provided their written informed consent. Participants completed two consecutive 45-minute scans, separated by brief (∼5-10 minutes) removal from the scanner. All scans were conducted on the same 3.0 Tesla Philips Ingenia Elition MRI scanner (Philips Healthcare, The Netherlands) using a 32-channel head coil. For voxel positioning, we first collected a *T*_1_-weighted structural MRI scan using the following parameters: compressed SENSE, TR/TE = 2 ms/2 ms, flip angle = 8°, slice thickness = 1.0 mm, 170 slices, voxel size = 1 mm^3^ isotropic, total time = 2 min 19 sec. Next, we acquired metabolite spectra from a 30 × 30 × 30 mm^3^ voxel in the bilateral dorsal anterior cingulate cortex (dACC; Figure 1). We positioned the dACC voxel on the mid-sagittal slice, just superior to the genu of the corpus callosum.**t**

**Fig 1.**
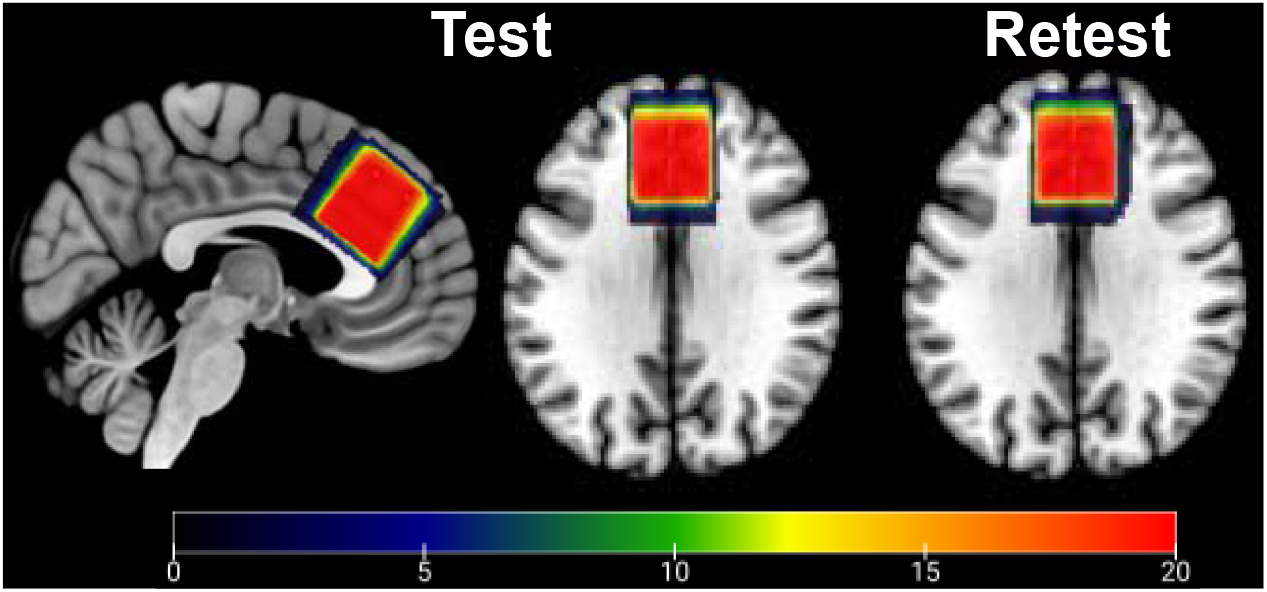
Voxel Placement. MR spectra were acquired from the dorsal anterior cingulate cortex (dACC); each participant’s native space binary voxel mask for their test (left) and retest (right) scans was normalized to standard (MNI) space and overlaid onto the spm152 template. Warmer colors indicate areas of greater overlap between participants (color bar = number of subjects overlapped).

We collected four consecutive metabolite spectra per scan (i.e., 4 “ test” and 4 “ retest” spectra per participant): two GABA-edited MEGA-PRESS sequences (i.e., with 14 or 20 ms editing pulses) and two HERMES sequences (i.e., with 14 or 20 ms editing pulses, each of which edited for both GABA and GSH). The order of sequences was randomized and balanced across subjects. Common parameters for the four sequences included: TR/TE = 2000 ms/80 ms; 224 averages in total; and MOIST water suppression (140 Hz bandwidth). The MEGA-PRESS sequences applied editing pulses at 1.9 ppm and 7.46 ppm; the HERMES sequences applied editing pulse lobes at 1.9 ppm and 4.56 ppm. Editing pulse duration was either 14 ms (referred to here as MEGA-14 or HERMES-14) or 20 ms (referred to here as MEGA-20 or HERMES-20). IWR was set to “ ON” for 10 participants and to “ OFF” for the remaining 10 participants (for both their test and retest acquisitions).

### 2.2 MRS Data Processing

MRS data were analyzed using the open-source analysis toolbox Osprey (v2.4.0; https://github.com/schorschinho/osprey/)^26^ within MATLAB R2021b. All analysis procedures followed consensus-recommended guidelines^19,29,30^. Briefly, analysis steps included: loading the vendor-native raw data; removing the residual water signal using a Hankel singular value decomposition (HSVD) filter^31^; eddy-current correction based on the water reference^32^; and robust spectral registration^33^ to align the individual transients within each sub-spectrum set (edit-ON and edit-OFF for MEGA-PRESS and sub-spectrum A, B, C, and D for HERMES). Final alignment of the averaged MEGA-PRESS sub-spectra minimized the residual water peak in the difference spectrum before subtraction to generate the GABA-edited difference spectrum. The HERMES sub-spectra were aligned using three pairwise steps, adjusting the frequency and phase such that the residual water was minimized for the GABA-ON/GSH-OFF sub-spectrum (aligning B to D), the 2 ppm tNAA signal was minimized to align the GABA-OFF/GSH-ON sub-spectrum (aligning C to D), and the 3.2 ppm tCho peak was minimized to align the GABA-ON/GSH-ON sub-spectrum (aligning A to C). The final GABA- and GSH-edited HERMES difference spectra were generated via Hadamard combination.

We modeled the metabolite spectra as detailed in our prior work^26,28,34^, using a wide modeling range (0.5-4 ppm^28^), either 0.4 or 0.55 ppm knot spacing, and custom basis sets for each sequence generated by our MRSCloud tool (https://braingps.mricloud.org/mrs-cloud^35^). We generated a basis set for each of the four sequences using PRESS localization at TE = 80 ms with the sequence-specific timing and real RF pulse waveforms and 14- or 20-ms editing pulse durations. The basis sets consisted of 19 basis functions (ascorbate, aspartate, creatine (Cr), negative creatine methylene (-CrCH_2_), GABA, glycerophosphocholine, glutathione (GSH), glutamine, glutamate, H_2_O, lactate, myo-inositol, NAA, NAAG, phosphocholine, phosphocreatine, phosphoethanolamine, scyllo-inositol, and taurine). For the edit-OFF (MEGA-PRESS) and the sum spectrum (HERMES), 5 macromolecules (MM09, MM12, MM14, MM17) and 3 lipids (Lip09, Lip13, Lip20) were included as parameterized Gaussian basis functions (see Zöllner et al., 2021 for details). The GABA-edited difference spectrum included parameterized Gaussian basis functions for the 0.9 and 3 ppm macromolecule signals (Zöllner et al, 2022). We modeled the co-edited MMs at 3 ppm using three different MM models^28^:

- **1to1GABA:** uses one single composite GABA + MM basis function by adding the edited GABA at 3 ppm and edited MM at 3 ppm (MM_3co_) with a fixed 1:1 amplitude ratio. That is, the 1to1GABA model assumes that 50% of the 3 ppm GABA signal is attributed to coedited MM.
- **1to1GABAsoft:** instead of using a hard 1:1 assumption for the relationship between GABA and MM_3co_, this model uses separate GABA and MM_3co_ basis functions and imposes a soft 1:1 amplitude constraint on the ratio of the GABA and MM_3co_ basis functions during the optimization step.
- **3to2MM:** uses separate basis functions for GABA and MM. The MM basis function in this model comprises both the MMs at 0.94 ppm (MM_0.94_) and the MMs at 3 ppm (MM_3co_), added in a 3:2 ratio. This model does not impose any amplitude assumptions or constraints on GABA.

Note that, although we model the MM signal in these ways, the primary outcome variable that we report is still “ GABA+” (GABA+MM), as is standard for 3 T GABA-edited MEGA-PRESS data. This is necessary to compare the three MM models, as different levels of constraint on the GABA:MM ratio (ranging from none for 3to2MM to full in 1to1GABA) change the covariance between GABA and MM. Modeling of MM in some fashion is necessary to appropriately model GABA+-edited spectra, but the inclusion of two covarying components in the model does not assume that GABA and MM can be meaningfully resolved with this acquisition methodology.

Binary masks of the test and retest MRS voxels were reconstructed in subject space and coregistered to each participant’s corresponding test or retest *T*_1_-weighted structural scan. We then segmented the structural scans using SPM12^36^ and quantified metabolite levels with respect to the unsuppressed water scan. In order to investigate variance inherent to MRS (as opposed to additional variance introduced by factors such as tissue segmentation), similar to our prior test-retest work^14^, no further relaxation or tissue segmentation corrections were applied. For additional details, see Appendix A, in which we list all consensus-recommended parameters regarding our MRS data acquisition, processing, and quality^37^.

We also processed all MRS data using Gannet^27^; see Appendix B3 for details. We present comparisons between within-subject reproducibility of metabolite concentrations generated by Osprey versus Gannet as a supplemental analysis. Gannet analyses are presented with the intention of replicating our previous results showing better within-subject reproducibility using Osprey’s linear-combination modeling procedures compared with Gannet’s simple peak fitting approach^14^.

### 2.3 Statistical Analyses

We conducted all statistical analyses using R 4.0.0^38^ within RStudio^39^. First, as in a majority of previous GABA-edited MRS test-retest work^11,12,14,15,40^, we estimated the between-scan intrasubject reproducibility of GABA+ estimates from each acquisition and modeling condition using within-subject CVs. These CVs provide a metric of within-subject measurement agreement that is independent of the range of values in the sample. We calculated within-subject CVs and confidence intervals using the root-mean method according to (Bland, 2006: https://www-users.york.ac.uk/~mb55/meas/cv.htm); the R function for this calculation is available at https://github.com/khupfeld/within-subject-cv. We focus the Results section on the impact of acquisition and analysis on within-subject CVs (the most commonly reported test-retest repeatability metric in the MRS literature). However, for completeness, in Appendix B1 we also report in two other common reliability metrics in the MRS test-retest literature^11^: Pearson correlation (i.e., the direction and strength of relationship between the test and retest values) and intraclass correlation coefficient (ICC, a metric of how reliably an instrument distinguishes between subjects, accounting for both the consistency of within-subject values between test and retest, as well as the change in group-average values between test and retest^41^). Single-rater, absolute-agreement, two-way mixed effect ICCs^11,42^ were calculated using the *irr* package^43^.

## 3. Results

### 3.1 GABA+ Data Quality

Three out of 80 total test-retest datasets (one MEGA-14 dataset and two HERMES-20 datasets) were excluded before any statistical analyses due to incorrect editing pulse parameters, unacceptably large lipid signal, and failure of sub-spectra alignment during post-processing. Thus, data presented in the Results include MEGA-14 (n=19), MEGA-20 (n=20), HERMES-14 (n=20), and HERMES-20 (n=18).

Group mean spectra for each sequence are presented in Figure 2. Creatine (Cr) linewidths (overall mean 5.4 Hz) were well within the range of consensus-recommended standards, i.e., < 13 Hz for 3T^30^ and did not differ between test and retest for any sequence, indicating consistent data quality across conditions (Table 1). Gray- and white-matter tissue fractions were also highly correlated between test and retest (fGM: Pearson r = 0.93, *p* < 0.001; fWM: Pearson r = 0.95, *p* < 0.001) and did not differ between test and retest for any sequence (fGM: t = 1.22; *p* = 0.236; fWM: t = −0.10; *p* = 0.923), indicating reproducible tissue inclusion across conditions. Additional consensus-recommended data quality metrics are presented in Appendix A.

**Table 1.** Data quality for test and retest conditions for each sequence

| Sequence | Test<br>Mean (SD) | Retest<br>Mean (SD) | t | p |
| --- | --- | --- | --- | --- |
| **MEGA-14** |  |  |  |  |
| Cr Linewidth (Hz) | 5.37 (0.71) | 5.40 (0.64) | -0.16 | 0.875 |
| Cr SNR<sup>a</sup> | 135.04 (23.86) | 130.59 (21.93) | 1.12 | 0.278 |
| **MEGA-20** |  |  |  |  |
| Cr Linewidth (Hz) | 5.35 (0.64) | 5.29 (0.60) | 0.38 | 0.707 |
| Cr SNR | 134.69 (25.50) | 131.39 (22.41) | 0.88 | 0.389 |
| **HERMES-14** |  |  |  |  |
| Cr Linewidth (Hz) | 5.58 (0.79) | 5.56 (0.73) | 0.09 | 0.932 |
| Cr SNR | 192.21 (31.49) | 183.45 (31.95) | 1.41 | 0.175 |
| **HERMES-20** |  |  |  |  |
| Cr Linewidth (Hz) | 5.51 (0.76) | 5.45 (0.66) | 0.31 | 0.758 |
| Cr SNR | 190.84 (25.21) | 191.01 (30.69) | -0.03 | 0.975 |

*Note.* Mean (standard deviation (SD)) and results of a within-subjects paired t-test are presented for each quality metric. Cr SNR is the maximum amplitude of the tCr peak divided by the detrended standard deviation of the noise.
<sup>a</sup> In the one instance where the paired t-test normality assumption was not met, we also conducted a paired samples Wilcoxon signed rank test. The statistical significance of the Wilcoxon test did not differ from that of the paired t-test.

**Fig 2.**
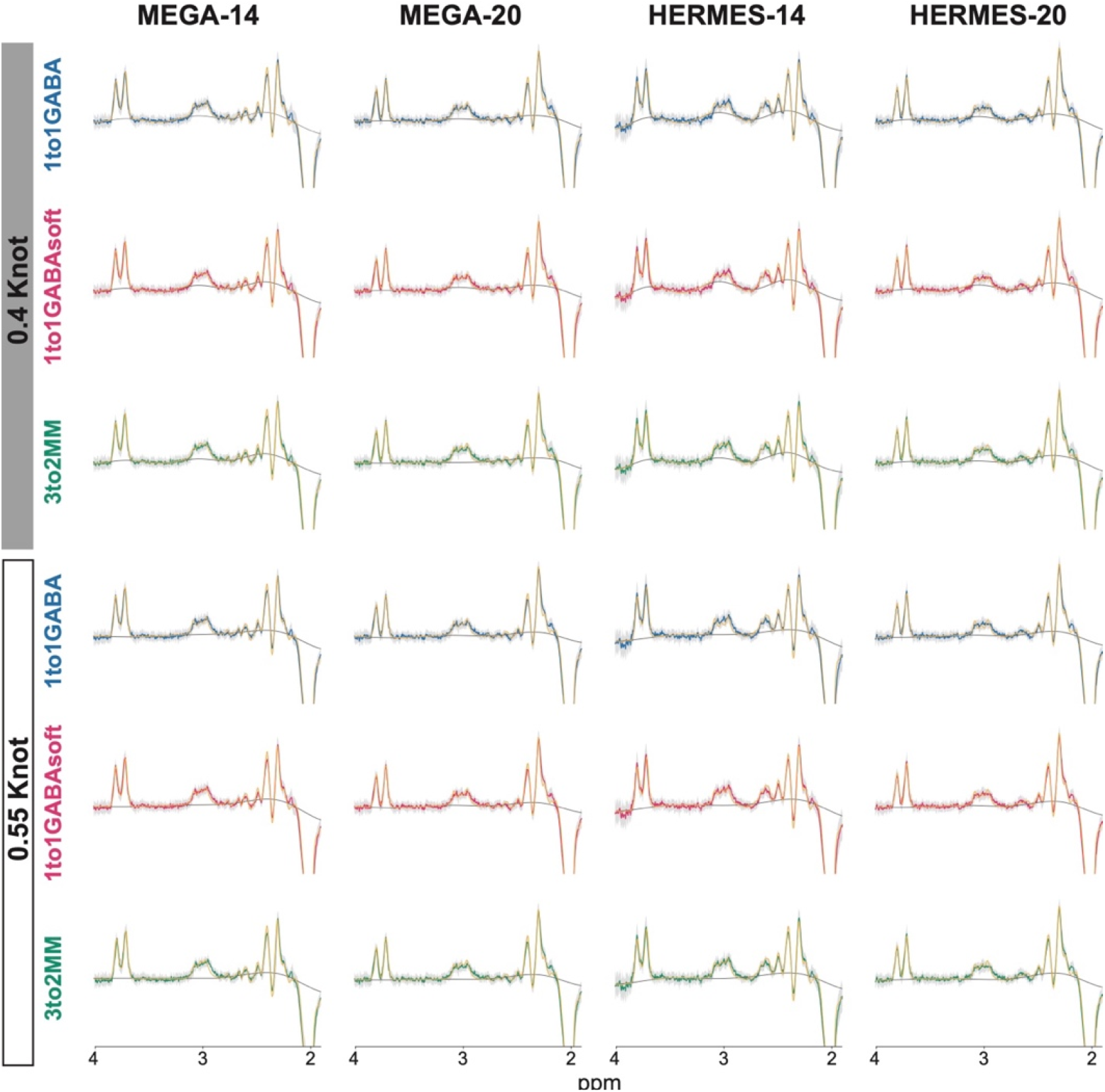
Group Mean GABA+ Spectra. Group average GABA-edited MEGA-PRESS and HERMES spectra (blue, pink, or green), 95% confidence interval (gray shading) and model fits (yellow) are shown for all participants for the test condition. Spectra are shown from 1.9-4.0 ppm.

### 3.2 GABA+ Within-Subject Reproducibility

Figure 3 depicts test and retest GABA+ values for each participant for each condition. Table 2 and Figure 4 indicate within-subject CVs and CV confidence intervals for GABA+ for each sequence and post-processing modeling option. Overall, there was not a consistent impact of acquisition sequence or editing pulse duration on GABA+ within-subject CVs, though HERMES CVs were generally more variable across acquisition and modeling choices than were MEGA CVs: MEGA-14 (mean 13.4%, range 8.1-21.0%), MEGA-20 (mean 12.4%, range 9.1-18.6%), HERMES-14 (mean 22.4%, range 11.1-34.4%), HERMES-20 (mean 13.4%, range 5.9-18.0%). Though HERMES-14 generally yielded the poorest reproducibility of the four acquisitions, this was the case only when the HERMES-14 data were fit using the 1to1GABA (29.2%, 34.4%) and 1to1GABAsoft (18.8%, 29.9%) models and not when using the 3to2MM model (11.1%, 11.3%).

**Table 2.** Within-subject CVs for GABA+

|  |  | MEGA-14 | MEGA-20 | HERMES-14 | HERMES-20 |
| --- | --- | --- | --- | --- | --- |
| 0.4 ppm knot | 1to1GABA | **21.0%**<br>(11.7-27.2%) | **15.3%**<br>(12.4-17.8%) | **34.4%**<br>(26.0-41.1%) | **18.0%**<br>(11.1-23.0%) |
|  | 1to1GABAsoft | **16.6%**<br>(8.1-22.1%) | **18.6%**<br>(11.7-23.6%) | **29.9%**<br>(19.2-37.7%) | **17.2%**<br>(9.8-22.2%) |
|  | 3to2MM | **16.7%**<br>(11.6-20.6%) | **9.9%**<br>(8.2-11.3%) | **11.3%**<br>(7.4-14.2%) | **11.4%**<br>(9.0-13.4%) |
| 0.55 ppm knot | 1to1GABA | **8.1%**<br>(2.6-11.2%) | **9.1%**<br>(4.6-12.0%) | **29.2%**<br>(23.5-33.9%) | **15.3%**<br>(7.7-20.2%) |
|  | 1to1GABAsoft | **9.1%**<br>(6.5-11.1%) | **11.2%**<br>(7.9-13.7%) | **18.8%**<br>(14.0-22.6%) | **12.7%**<br>(7.8-16.2%) |
|  | 3to2MM | **9.1%**<br>(5.1-11.8%) | **10.0%**<br>(7.8-11.8%) | **11.1%**<br>(6.8-14.1%) | **5.9%**<br>(4.6-6.9%) |

Note. Within-subject coefficients of variation (CVs) and 90% confidence intervals (CIs) were calculated according to the root mean square method described by (Bland, 2006: https://www-users.york.ac.uk/~mb55/meas/cv.htm).

**Fig 3.**
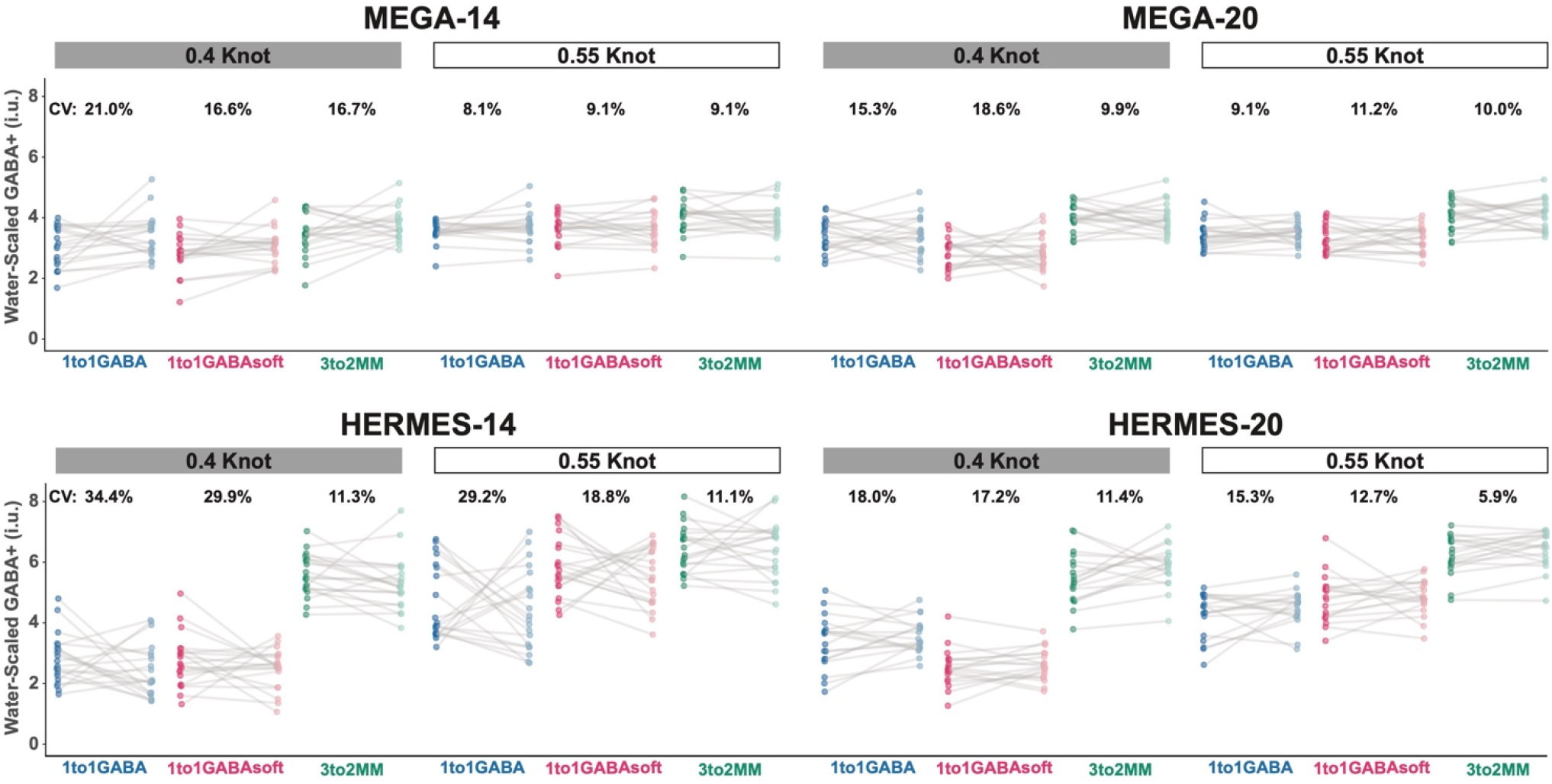
Within-Subject GABA+ values. GABA+ values are shown for the test (dark) and retest (light) conditions. Each point and connecting gray line represents one participant. Within-subject CV percentages are listed above each subplot.

**Fig 4.**
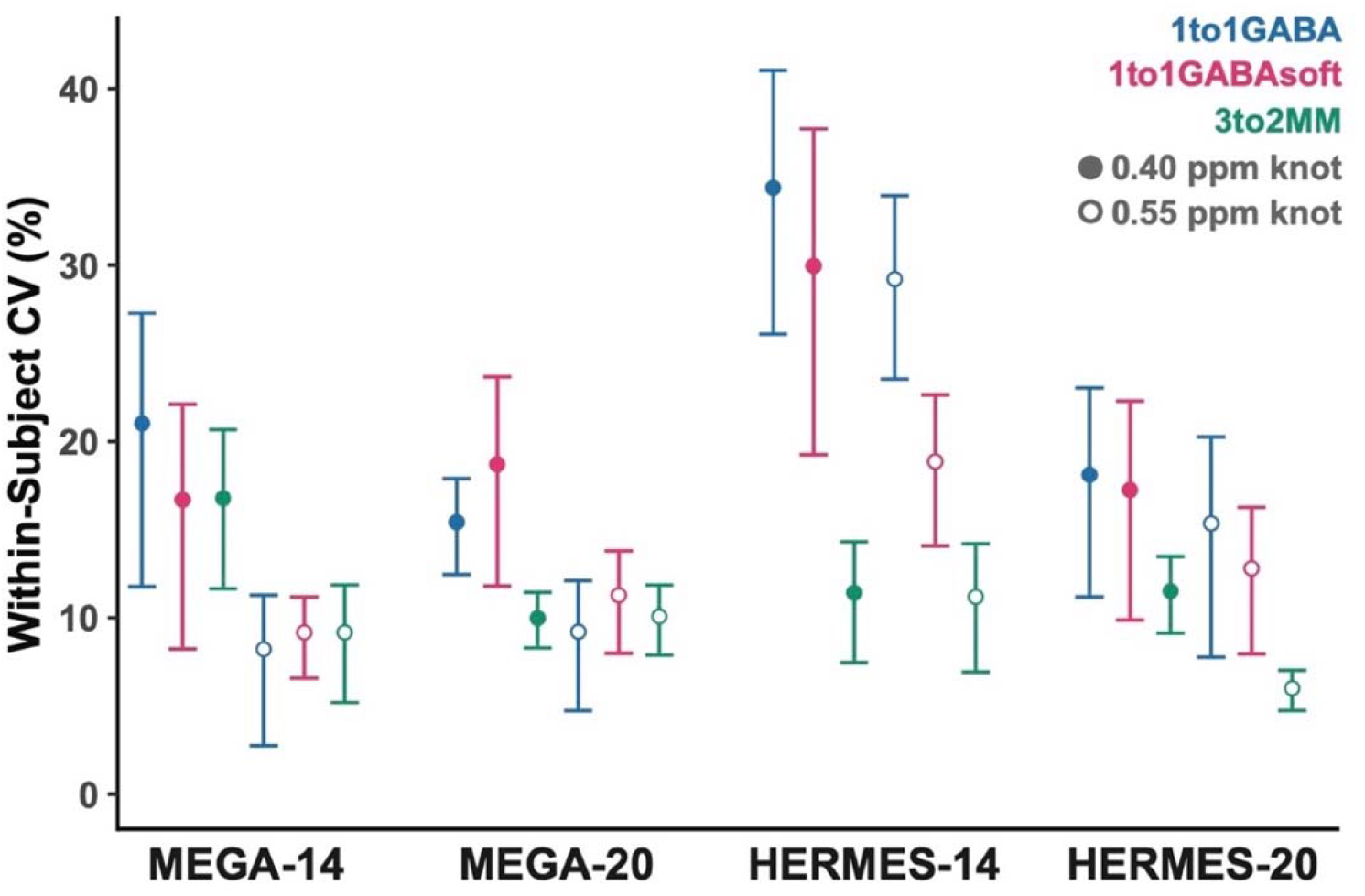
Within-subject CVs for GABA+. Within-subject %CVs and 90% confidence intervals are shown for each acquisition and post-processing condition.

Indeed, across all four experiments, the 3to2MM model generally yielded the best reproducibility (mean 10.7%, range 5.9-16.7%) compared with the 1to1GABA (mean 18.8%, range 8.1-34.4%) and 1to1GABAsoft models (mean 16.8%, range 9.1-29.9%). This benefit of the 3to2MM model was particularly evident for the HERMES data. Across all conditions, 0.55 ppm spacing generally performed better (mean 12.5%, range 5.9-29.2%) than 0.4 ppm knot spacing (mean 18.4%, range 9.9-34.4%). The benefit of sparser knot spacing was somewhat larger for the MEGA-PRESS data (0.55 ppm mean 9.4%; 0.4 ppm mean 16.4%) compared with the HERMES data (0.55 ppm mean 15.5%; 0.4 ppm mean 20.4%). ICC values and Pearson correlation coefficients generally corresponded with within-subject CVs and provide further support for these conclusions (Tables B1.1-B1.2).

We encountered minimal drift in either IWR condition (IWR-ON: 0.029 ± 0.004 ppm; IWR-OFF: 0.040 ± 0.016 ppm) and drift did not differ between the IWR-ON versus IWR-OFF or test versus retest conditions (both *p* > 0.05; Table B2.1). For completeness, we present within-subject CVs by IWR conditions in Table B2.2. With the exception of MEGA-14, CV values were generally slightly better (lower) for IWR-ON compared with IWR-OFF: MEGA-14 (mean-ON 13.5%, range-ON 5.7-22.9%; mean-OFF 13.2%, range-OFF 9.4-19.0%), MEGA-20 (mean-ON 9.6%, range-ON 7.2-13.2%; mean-OFF 14.4%, range-OFF 9.7-24.7%), HERMES-14 (mean-ON 20.7%, range-ON 8.4-31.9%; mean-OFF 24.0%, range-OFF 11.8-36.7%), and HERMES-20 (mean-ON 11.9%, range-ON 5.8-16.4%; mean-OFF 14.8%, range-OFF 6.0-22.6%). However, given the lack of significant difference in scanner stability between IWR conditions, these results should be interpreted with caution.

### 3.3 GABA+ Results Using Gannet Modeling

For completeness and comparison with prior literature, Gannet modeling results are presented in Appendix B3. For each acquisition within-subject CVs for GABA+ were higher (indicating poorer test-retest reproducibility) for Gannet compared with Osprey. This effect was strongest for 20 ms editing pulses: MEGA-14 (mean Osprey CV 13.4%, Gannet CV 17.7%), MEGA-20 (mean Osprey CV 12.4%, Gannet CV 25.4%), HERMES-14 (mean Osprey CV 22.5%, Gannet CV 25.5%), and HERMES-20 (mean Osprey CV 13.4%, Gannet CV 28.6%).

### 3.4 GSH Within-Subject Reproducibility

For completeness, we followed the same procedures to calculate within-subject CVs for GSH measured with the HERMES-14 and HERMES-20 sequences (Table B4.1; Figures B4.3-B4.4). Editing pulse duration did not have a clear impact on GSH CV: HERMES-14 (mean 15.7%) and HERMES-20 (mean 15.1%). The co-edited MM models apply only to GABA+ and not GSH and thus only spline baseline knot spacing was investigated in relation to GSH. Contrary to the case for GABA+, less sparse spline baseline knot spacing resulted in lower CVs for GSH, though this effect was more pronounced for HERMES-20 (0.4 ppm 12.9%, 0.55 ppm 17.3%) compared with HERMES-14 (0.4 ppm 15.2%, 0.55 ppm 16.2%). Similar to GABA+, within-subject CVs for GSH were worse following Gannet compared with Osprey for both HERMES-14 (mean Osprey CV 15.7%, Gannet CV 36.1%) and HERMES-20 (mean Osprey CV 12.9%, Gannet CV 41.2%).

## 4. Discussion

While sequence complexity and editing pulse duration did not consistently affect GABA+ reproducibility, we identified the overall best test-retest reproducibility (lowest within-subject CVs) following post-processing modeling of co-edited MMs using the 3to2MM model and using sparser (0.55 ppm) spline baseline knot spacing. The 3to2MM model particularly benefited the HERMES data; 0.55 ppm baseline knot spacing benefited MEGA-PRESS somewhat more than HERMES data. Linear-combination modeling with Osprey resulted in better reproducibility for GABA+ than the simple peak fitting approach implemented in Gannet.

Overall, the more complex, 4-experiment HERMES sequence did not result in consistently different test-retest reproducibility of GABA+ compared with the 2-experiment MEGA-PRESS sequence. We anticipated HERMES CVs to be worse than MEGA-PRESS CVs for multiple reasons; for example, HERMES requires successful alignment of more sub-spectra and thus is more susceptible to artifacts from subject movement. Similarly, editing pulse duration did not consistently influence GABA+ reproducibility. We anticipated that shorter, less selective 14 ms editing pulses would result in less susceptibility to motion and scanner drift artifacts^19^ and result in better GABA+ reproducibility. However, given that this was a healthy control cohort, we did not encounter significant motion in the participants. Moreover, we did not have significant frequency drift; thus, susceptibility to drift likely did not have a substantial effect on this cohort (though drift could have more influence in a patient or pediatric cohort or for scans acquired on a different scanner or during an EPI-heavy protocol).

HERMES-14 generally yielded the poorest reproducibility of the four acquisitions. HERMES-14 is an experiment that was included in this protocol to complete the 2×2 design, without any further development or optimization. It is thus perhaps not surprising that is performed poorly, as we had previously assumed that HERMES would not work with 14 ms pulses. However, both HERMES-14 and HERMES-20 reproducibility were poorest when the data were fit using the 1to1GABA and 1to1GABAsoft models compared with the 3to2MM model. HERMES CVs were also consistently more variable than were MEGA-PRESS CVs. It thus appears that modeling choices may exert a greater influence on reproducibility of HERMES GABA+ data than MEGA-PRESS data, and that the 3to2MM model may fit HERMES GABA+ data most consistently—though it remains unclear why specifically this is the case.

In addition to yielding the best reproducibility for HERMES GABA+, the 3to2MM model generally resulted in the smallest GABA+ CVs (best reproducibility) across all acquisitions. The 3to2MM model adds separate basis functions for GABA+ and MM (both MM_0.94_ and MM_3co_) in a 3:2 ratio and does not impose amplitude constraints on GABA+. Benefits of the 3to2MM model include the addition of the MM_0.94_ peak, which provides a non-overlapped anchoring reference for the amplitude of the MM_3co_ peak and exploits the fact that MM profiles are relatively consistent across healthy participants and ages^44^. In our prior work in MEGA-PRESS TE = 68 ms GABA+ data^28^, we found some of the lowest intersubject CVs using the 3to2MM model; this model performed well without overfitting the data. Though we hesitate to make any definitive recommendations regarding the optimal modeling decisions from the current sample size, these TE = 80 ms GABA+ data suggest that 3to2MM is likely a good model to use for GABA+ across a variety of acquisition parameters, at least in the absence of pathological MM changes. It is not easy to say which model is more valid—our ability to estimate the correct GABA:MM ratio determines our ability to estimate the correct MM_3.0_:MM_0.9_ ratio. It is notable that the most reproducible model (3to2MM) has the greatest freedom (the GABA and MM components are not relatively constrained) to model signal at 3 ppm. It is also interesting that the 1to1GABA constraints perform poorly across both 14 and 20 ms editing pulse durations, as the relative contribution of MM to the GABA signal is lower (and more variable) for the more selective (20 ms) editing pulses.

Sparser 0.55 ppm spline baseline knot spacing yielded better GABA+ reproducibility overall, and particularly for MEGA-PRESS GABA+. This is similar to our prior report that 0.55 ppm knot spacing resulted in lower intersubject CVs for TE = 68 ms GABA+ data^28^. It is logical that the sparser (i.e., more rigid) baseline better estimated GABA+, as the more rigid baseline did not tend to overfit the signal, i.e., did not incorrectly assign some of the GABA+ signal to baseline (as was the case for the less sparse 0.4 ppm baseline which tended to bend upwards into the GABA+ signal). Contrarily, we found that more flexible 0.4 ppm knot spacing yielded better reproducibility for GSH. This effect was particularly evident for HERMES-20 compared with HERMES-14 GSH; this finding will require further investigation but likely allows the model to better accommodate the less flat baseline behavior in GSH-edited spectra. The results here highlight primarily that spline baseline knot spacing decisions can greatly affect reproducibility of metabolite estimates and that this modeling decision should likely differ depending on the metabolite of interest for a given study.

We found better within-subject reproducibility for both GABA+ and GSH following linear-combination modeling in Osprey compared with simple peak fitting in Gannet. This is similar to our prior test-retest work^14^, which reported significant improvement in HERMES GSH test-retest reproducibility with Osprey (CV 9%) over Gannet (CV 19%). In the present work, we found this CV improvement with Osprey for both GABA+ and GSH and for both HERMES and MEGA-PRESS, albeit with worse overall reproducibility. In Song et al.^14^, we saw comparable GABA+ CVs to the present work but did not see significant improvement in HERMES GABA+ CVs between the two modeling approaches (Osprey CV 15%, Gannet CV 17%)--though this prior work used only the 1to1GABAsoft model and Osprey default 0.4 ppm spline baseline knot spacing, which was found here to perform worse for GABA+. While there were acquisition differences between the two studies (e.g., different scanner vendor) it remains unclear precisely why GSH reproducibility was overall worse in the present work compared with Song et al. The ability of low-n test-retest studies to truly characterize reproducibility is limited. Nonetheless, our prior and present results do tend to support implementing consensus-recommended linear combination modeling^29^ for quantifying edited metabolites collected using various acquisitions.

There are several limitations to this work. The dACC is a moderately challenging region for MRS which is more susceptible to artifacts and low SNR than e.g., posterior cingulate cortex; however, we selected this region because of its clinical relevance to cognition^45^ and to match with our prior test-retest work^12,14^. Though (as anticipated) IWR-ON resulted in slightly better reproducibility compared with IWR-OFF, we did not encounter substantial scanner frequency drift in this dataset. Benefits of using IWR would likely become more evident for within-subject measurements taken over multiple days, on scanners with greater drift, or in patient cohorts with greater movement. There is no gold standard of metabolite level estimation for GABA+ to validate the results. Though the performance of MRS acquisition and modeling approaches is often judged by the amount of within-subject variance^11,12,14,15,40^, lower variance does not necessarily reflect greater accuracy (but is one pre-condition for it). However, as these healthy control participants did not undergo any intervention between the test and retest phases of the experiment, CVs should predominantly reflect variance introduced by the acquisition and modeling parameters. Finally, this was a limited-sample-size dataset and we therefore focus our interpretations on differences in descriptive statistics (primarily, differences in within-subject CVs). We designed this experiment with the intention of comparing acquisition parameters, but *post hoc* decided to additionally explore the impact of modeling decisions on test-retest reproducibility. Thus, given the sample size, we do not suggest that our findings should be interpreted as definitive recommendations for optimal GABA+ acquisition or modeling (although they are perhaps the best evidence available on which to make such decisions). Rather, we intend to highlight that reproducibility differs based on acquisition and modeling parameters and that such parameters could significantly affect interpretation of results in clinical work. Future studies in larger datasets should be conducted to provide more concrete recommendations for acquisition and modeling of edited GABA+ data at TE = 80 ms.

## Ethics Approval and Consent to Participate

The Johns Hopkins University Institutional Review Board approved all study procedures, and written informed consent was obtained from all participants.

## Consent for Publication

All authors consent to the publication of this study.

## Availability of Data and Material

The raw data supporting the conclusions of this manuscript will be made available by the authors without undue reservation.

## Author Contributions

KH collected and processed the data, conducted all statistical analyses, created all figures and supplemental material, and wrote the first draft of the manuscript. HZ and GO advised on MRS processing. YS assisted with data collection. SH conducted all simulations. VY reviewed all structural scans to assess data quality and check for incidental findings. JP and RAE designed the project and led interpretation of the results. All authors participated in revision of the manuscript.

