## Appendix A for "Impact of acquisition and modeling parameters on test-retest reproducibility of edited GABA+"

### Appendix A: Summary MRSinMRS Report

Here we provide a summary following the minimum reporting standards in MRS generated in Osprey. For further details, see: Lin et al. 'Minimum Reporting Standards for *in vivo* Magnetic Resonance Spectroscopy (MRSinMRS): Experts' consensus recommendations. NMR in Biomedicine. 2021;e4484. doi.org/10.1002/nbm.4448

**Site:** The Johns Hopkins University School of Medicine / F. M. Kirby Research Center for Functional Brain Imaging, Kennedy Krieger Institute

| **1. Hardware** | |
|---|---|
| a. Field strength [T] | 3 T |
| b. Manufacturer | Philips |
| c. Model (software version if available) | R 5.1.7 |
| d. RF coils: nuclei (transmit/receive), number of channels, type, body part | 1H, 32 channel, head |
| e. Additional hardware | - |

| **2. Acquisition** | |
|---|---|
| **MEGA-14 and MEGA-20** | |
| a. Pulse sequence | MEGA-PRESS (Johns Hopkins University Patch) |
| b. Volume of interest (VOI) location | Dorsal ACC |
| c. Nominal VOI size [$mm^3$] | 30 x 30 x 30 $mm^3$ |
| d. Repetition time (TR), echo time (TE) [ms] | TR 2000 ms, TE 80 ms |
| e. Total number of averages per spectrum | 224 total averages |
| i. Number of averaged spectra per subspectrum | with 112 averages per subspectrum |
| f. Additional sequence parameters | F1: 2000 Hz, 2048 points |
| i. editing pulse frequencies | $ppm_{ON}$ = 1.90, $ppm_{OFF}$ = 7.46 |
| g. Water suppression method | MOIST |
| h. Shimming method, reference peak, and threshold of acceptance of shim chosen | 2nd order pencil beam, water |
| i. Trigger or motion correction | No trigger or active motion correction |
| **HERMES-14 and HERMES-20** | |
| a. Pulse sequence | HERMES (Johns Hopkins University Patch) |
| b. Volume of interest (VOI) location | Dorsal ACC |
| c. Nominal VOI size [$mm^3$] | 30 x 30 x 30 $mm^3$ |
| d. Repetition time (TR), echo time (TE) [ms] | TR 2000 ms, TE 80 ms |
| e. Total number of averages per spectrum | 224 total averages |
| i. Number of averaged spectra per subspectrum | with 56 averages per subspectrum |
| f. Additional sequence parameters | F1: 2000 Hz, 2048 points |
| i. editing pulse frequencies | $ppm_{ON}$ = 1.90, $ppm_{OFF}$ = 4.56 |
| g. Water suppression method | MOIST |
| h. Shimming method, reference peak, and threshold of acceptance of shim chosen | 2nd order pencil beam, water |
| i. Trigger or motion correction | No trigger or active motion correction |

| 3. Data analysis methods and outputs | |
|---|---|
| **MEGA-14, MEGA-20, HERMES-14, and HERMES-20** | |
| a. Analysis software | Osprey 2.4.0 |
| b. Processing steps deviating from Osprey | None |
| c. Output measure | rawWaterScaled |
| d. Quantification references and assumptions, fitting model assumptions | Basis set list: Asc, Asp, Cr, -$CrCH_2$, GABA, GPC, GSH, Gln, Glu, $H_2O$, Lac, mI, NAA, NAAG, PCh, PCr, PE, sI, Tau. Edit-OFF (MEGA) and Sum (HERMES): MM09, MM12, MM14, MM17, Lip09, Lip13, Lip20; GABA-edited DIFF (MEGA and HERMES): $MM_{3co}$ and $MM_{0.94}$. Fitting method: Osprey baseline knot spacing 0.40 ppm or 0.55 ppm |

| 4. Data quality | |
|---|---|
| **MEGA-14-Test** | |
| a. SNR (Cr), linewidth (Cr) [Hz, OFF spectra] | SNR: 135 ± 24, linewidth: 5.37 ± 0.71 Hz |
| b. Data exclusion criteria | Cr linewidth > 13 Hz |
| c. Quality measures of postprocessing model fitting (Mean Relative Amplitude Residual) | |
| ***0.4 ppm, 1to1GABA*** | |
| OFF | 13.08 ± 8.81 |
| DIFF1 | 2.97 ± 0.93 |
| ***0.4 ppm, 1to1GABAsoft*** | |
| OFF | 13.08 ± 8.81 |
| DIFF1 | 2.75 ± 0.66 |
| ***0.4 ppm, 3to2MM*** | |
| OFF | 13.08 ± 8.81 |
| DIFF1 | 2.71 ± 0.61 |
| ***0.55 ppm, 1to1GABA*** | |
| OFF | 15.95 ± 10.32 |
| DIFF1 | 2.71 ± 0.60 |
| ***0.55 ppm, 1to1GABAsoft*** | |
| OFF | 15.95 ± 10.32 |
| DIFF1 | 2.95 ± 0.79 |
| ***0.55 ppm, 3to2MM*** | |
| OFF | 15.95 ± 10.32 |
| DIFF1 | 2.80 ± 0.64 |
| d. Mean spectrum created with OspreyOverview | Figure 2 |

| | |
|---|---|
| **4. Data quality** | |
| **MEGA-14-Retest** | |
| a. SNR (Cr), linewidth (Cr) [Hz, OFF spectra] | SNR: 131 ± 22, linewidth: 5.40 ± 0.64 Hz |
| b. Data exclusion criteria | Cr linewidth > 13 Hz |
| c. Quality measures of postprocessing model fitting (Mean Relative Amplitude Residual) | |
| ***0.4 ppm, 1to1GABA*** | |
| OFF | 11.15 ± 4.63 |
| DIFF1 | 2.65 ± 0.46 |
| ***0.4 ppm, 1to1GABAsoft*** | |
| OFF | 11.15 ± 4.63 |
| DIFF1 | 2.60 ± 0.54 |
| ***0.4 ppm, 3to2MM*** | |
| OFF | 11.15 ± 4.63 |
| DIFF1 | 2.55 ± 0.50 |
| ***0.55 ppm, 1to1GABA*** | |
| OFF | 13.54 ± 5.97 |
| DIFF1 | 2.81 ± 1.26 |
| ***0.55 ppm, 1to1GABAsoft*** | |
| OFF | 13.54 ± 5.97 |
| DIFF1 | 2.88 ± 1.23 |
| ***0.55 ppm, 3to2MM*** | |
| OFF | 13.54 ± 5.97 |
| DIFF1 | 2.98 ± 1.40 |
| d. Mean spectrum created with OspreyOverview | Figure 2 |
| **MEGA-20-Test** | |
| a. SNR (Cr), linewidth (Cr) [Hz, OFF spectra] | SNR: 135 ± 26, linewidth: 5.35 ± 0.64 Hz |
| b. Data exclusion criteria | Cr linewidth > 13 Hz |
| c. Quality measures of postprocessing model fitting (Mean Relative Amplitude Residual) | |
| ***0.4 ppm, 1to1GABA*** | |
| OFF | 13.56 ± 8.99 |
| DIFF1 | 2.75 ± 0.56 |
| ***0.4 ppm, 1to1GABAsoft*** | |
| OFF | 13.56 ± 8.99 |
| DIFF1 | 2.62 ± 0.49 |
| ***0.4 ppm, 3to2MM*** | |
| OFF | 13.56 ± 8.99 |
| DIFF1 | 2.64 ± 0.53 |
| ***0.55 ppm, 1to1GABA*** | |
| OFF | 16.10 ± 11.05 |
| DIFF1 | 2.56 ± 0.45 |
| ***0.55 ppm, 1to1GABAsoft*** | |
| OFF | 16.10 ± 11.05 |
| DIFF1 | 2.57 ± 0.49 |
| ***0.55 ppm, 3to2MM*** | |
| OFF | 16.10 ± 11.05 |
| DIFF1 | 2.77 ± 0.51 |
| d. Mean spectrum created with OspreyOverview | Figure 2 |

#### 4. Data quality

**MEGA-20-Retest**

| | |
|---|---|
| a. SNR (Cr), linewidth (Cr) [Hz, OFF spectra] | SNR: 131 ± 22, linewidth: 5.29 ± 0.60 Hz |
| b. Data exclusion criteria | Cr linewidth > 13 Hz |
| c. Quality measures of postprocessing model fitting (Mean Relative Amplitude Residual) | |
| ***0.4 ppm, 1to1GABA*** | |
| OFF | 10.39 ± 4.24 |
| DIFF1 | 2.76 ± 1.00 |
| ***0.4 ppm, 1to1GABAsoft*** | |
| OFF | 10.39 ± 4.24 |
| DIFF1 | 2.63 ± 0.98 |
| ***0.4 ppm, 3to2MM*** | |
| OFF | 10.39 ± 4.24 |
| DIFF1 | 2.63 ± 0.94 |
| ***0.55 ppm, 1to1GABA*** | |
| OFF | 12.41 ± 6.26 |
| DIFF1 | 2.55 ± 0.93 |
| ***0.55 ppm, 1to1GABAsoft*** | |
| OFF | 12.41 ± 6.26 |
| DIFF1 | 2.57 ± 0.89 |
| ***0.55 ppm, 3to2MM*** | |
| OFF | 12.41 ± 6.26 |
| DIFF1 | 2.74 ± 0.94 |
| d. Mean spectrum created with OspreyOverview | Figure 2 |

**HERMES-14-Test**

| | |
|---|---|
| a. SNR (Cr), linewidth (Cr) [Hz, OFF spectra] | SNR: 192 ± 31, linewidth: 5.58 ± 0.79 Hz |
| b. Data exclusion criteria | Cr linewidth > 13 Hz |
| c. Quality measures of postprocessing model fitting (Mean Relative Amplitude Residual) | |
| ***0.4 ppm, 1to1GABA*** | |
| OFF | 17.98 ± 13.35 |
| DIFF1 | 3.62 ± 0.88 |
| DIFF2 | 2.84 ± 0.68 |
| ***0.4 ppm, 1to1GABAsoft*** | |
| OFF | 17.98 ± 13.35 |
| DIFF1 | 3.58 ± 0.99 |
| DIFF2 | 2.84 ± 0.68 |
| ***0.4 ppm, 3to2MM*** | |
| OFF | 17.98 ± 13.35 |
| DIFF1 | 3.63 ± 0.91 |
| DIFF2 | 2.84 ± 0.68 |
| ***0.55 ppm, 1to1GABA*** | |
| OFF | 21.59 ± 14.58 |
| DIFF1 | 3.96 ± 0.95 |
| DIFF2 | 2.94 ± 0.69 |
| ***0.55 ppm, 1to1GABAsoft*** | |
| OFF | 21.59 ± 14.58 |
| DIFF1 | 3.69 ± 0.98 |
| DIFF2 | 2.94 ± 0.69 |
| ***0.55 ppm, 3to2MM*** | |
| OFF | 21.59 ± 14.58 |
| DIFF1 | 3.77 ± 0.86 |
| DIFF2 | 2.94 ± 0.69 |
| d. Mean spectrum created with OspreyOverview | Figure 2 |

#### 4. Data quality

**HERMES-14-Retest**

| | |
|---|---|
| a. SNR (Cr), linewidth (Cr) [Hz, OFF spectra] | SNR: 183 ± 32, linewidth: 5.56 ± 0.73 Hz |
| b. Data exclusion criteria | Cr linewidth > 13 Hz |
| c. Quality measures of postprocessing model fitting (Mean Relative Amplitude Residual) | |
| ***0.4 ppm, 1to1GABA*** | |
| OFF | 12.58 ± 4.83 |
| DIFF1 | 3.30 ± 0.71 |
| DIFF2 | 3.03 ± 0.65 |
| ***0.4 ppm, 1to1GABAsoft*** | |
| OFF | 12.58 ± 4.83 |
| DIFF1 | 3.17 ± 0.76 |
| DIFF2 | 3.03 ± 0.65 |
| ***0.4 ppm, 3to2MM*** | |
| OFF | 12.58 ± 4.83 |
| DIFF1 | 3.20 ± 0.69 |
| DIFF2 | 3.03 ± 0.65 |
| ***0.55 ppm, 1to1GABA*** | |
| OFF | 16.36 ± 9.08 |
| DIFF1 | 3.50 ± 0.83 |
| DIFF2 | 3.11 ± 0.66 |
| ***0.55 ppm, 1to1GABAsoft*** | |
| OFF | 16.36 ± 9.08 |
| DIFF1 | 3.49 ± 1.05 |
| DIFF2 | 3.11 ± 0.66 |
| ***0.55 ppm, 3to2MM*** | |
| OFF | 16.36 ± 9.08 |
| DIFF1 | 3.46 ± 0.93 |
| DIFF2 | 3.11 ± 0.66 |
| d. Mean spectrum created with OspreyOverview | Figure 2 |

**HERMES-20-Test**

| | |
|---|---|
| a. SNR (Cr), linewidth (Cr) [Hz, OFF spectra] | SNR: 191 ± 25, linewidth: 5.51 ± 0.76 Hz |
| b. Data exclusion criteria | Cr linewidth > 13 Hz |
| c. Quality measures of postprocessing model fitting (Mean Relative Amplitude Residual) | |
| ***0.4 ppm, 1to1GABA*** | |
| OFF | 17.54 ± 15.05 |
| DIFF1 | 2.51 ± 0.53 |
| DIFF2 | 2.20 ± 0.47 |
| ***0.4 ppm, 1to1GABAsoft*** | |
| OFF | 17.54 ± 15.05 |
| DIFF1 | 2.56 ± 0.68 |
| DIFF2 | 2.20 ± 0.47 |
| ***0.4 ppm, 3to2MM*** | |
| OFF | 17.54 ± 15.05 |
| DIFF1 | 2.62 ± 0.60 |
| DIFF2 | 2.20 ± 0.47 |
| ***0.55 ppm, 1to1GABA*** | |
| OFF | 20.64 ± 16.04 |
| DIFF1 | 2.61 ± 0.54 |
| DIFF2 | 2.21 ± 0.46 |
| ***0.55 ppm, 1to1GABAsoft*** | |
| OFF | 20.64 ± 16.04 |
| DIFF1 | 2.57 ± 0.52 |
| DIFF2 | 2.21 ± 0.46 |

| | |
|---|---|
| **4. Data quality** | |
| ***0.55 ppm, 3to2MM*** | |
| OFF | 20.64 ± 16.04 |
| DIFF1 | 2.61 ± 0.51 |
| DIFF2 | 2.21 ± 0.46 |
| d. Mean spectrum created with OspreyOverview | Figure 2 |
| **HERMES-20-Retest** | |
| a. SNR (Cr), linewidth (Cr) [Hz, OFF spectra] | SNR: 191 ± 30.69, linewidth: 5.45 ± 0.66 Hz |
| b. Data exclusion criteria | Cr linewidth > 13 Hz |
| c. Quality measures of postprocessing model fitting (Mean Relative Amplitude Residual) | |
| ***0.4 ppm, 1to1GABA*** | |
| OFF | 14.18 ± 7.33 |
| DIFF1 | 2.59 ± 1.07 |
| DIFF2 | 2.21 ± 0.48 |
| ***0.4 ppm, 1to1GABAsoft*** | |
| OFF | 14.18 ± 7.33 |
| DIFF1 | 2.54 ± 0.95 |
| DIFF2 | 2.21 ± 0.48 |
| ***0.4 ppm, 3to2MM*** | |
| OFF | 14.18 ± 7.33 |
| DIFF1 | 2.67 ± 0.93 |
| DIFF2 | 2.21 ± 0.48 |
| ***0.55 ppm, 1to1GABA*** | |
| OFF | 18.06 ± 10.05 |
| DIFF1 | 2.65 ± 0.95 |
| DIFF2 | 2.20 ± 0.51 |
| ***0.55 ppm, 1to1GABAsoft*** | |
| OFF | 18.06 ± 10.05 |
| DIFF1 | 2.62 ± 0.99 |
| DIFF2 | 2.20 ± 0.51 |
| ***0.55 ppm, 3to2MM*** | |
| OFF | 18.06 ± 10.05 |
| DIFF1 | 2.73 ± 0.93 |
| DIFF2 | 2.20 ± 0.51 |
| d. Mean spectrum created with OspreyOverview | Figure 2 |

*Note.* Cr SNR is the maximum amplitude of the tCr peak divided by the detrended standard deviation of the noise. Mean relative amplitude residual is indicative of postprocessing model fit (in which higher values indicate poorer model fit).
