## Appendix B for "Impact of acquisition and modeling parameters on test-retest reproducibility of edited GABA+"

### Appendix B. Supplemental Figures and Tables

#### <u>Section B1</u>. GABA+ ICC and Pearson Correlations

**Table B1.1.** ICC values for GABA+

| | | **MEGA-14** | **MEGA-20** | **HERMES-14** | **HERMES-20** |
|---|---|---|---|---|---|
| 0.4 ppm knot | 1to1GABA | 0.07 | 0.21 | -0.18 | 0.41 |
| | 1to1GABAsoft | 0.43 | -0.06 | -0.02 | **0.58** |
| | 3to2MM | 0.24 | 0.25 | 0.36 | 0.34 |
| 0.55 ppm knot | 1to1GABA | **0.51** | 0.29 | -0.25 | 0.25 |
| | 1to1GABAsoft | **0.62** | 0.26 | -0.22 | 0.17 |
| | 3to2MM | **0.54** | 0.32 | 0.27 | **0.68** |

*Note*. Single-rater, absolute-agreement, two-way mixed effect ICCs[1,2] were calculated using the *irr* package[3]. ICC values < 0.50 indicate poor reliability; ICC values between 0.50 and 0.75 indicate moderate reliability[2] and are bolded.

**Table B1.2.** Pearson correlation coefficients for GABA+

| | | **MEGA-14** | **MEGA-20** | **HERMES-14** | **HERMES-20** |
|---|---|---|---|---|---|
| 0.4 ppm knot | 1to1GABA | 0.07 | 0.20 | -0.19 | 0.46 |
| | 1to1GABAsoft | **0.46*** | -0.06 | -0.02 | **0.58*** |
| | 3to2MM | 0.27 | 0.25 | 0.38 | 0.38 |
| 0.55 ppm knot | 1to1GABA | **0.56*** | 0.28 | -0.24 | 0.26 |
| | 1to1GABAsoft | **0.61**** | 0.26 | -0.22 | 0.17 |
| | 3to2MM | **0.54*** | 0.31 | 0.26 | **0.70**** |

*Note*. Pearson correlation coefficients were calculated between test and retest GABA+ estimates. Statistically significant correlations are bolded; *$p < 0.05$, **$p < 0.01$.

### Section B2. Frequency Drift

**Table B2.1.** Frequency drift for test versus retest and IWR-ON versus IWR-OFF

| | **Test Mean (SD)** | **Retest Mean (SD)** | **V** | ***p*** |
|---|---|---|---|---|
| Drift (abs(ΔCr (ppm)) | 0.035 (0.014) | 0.034 (0.014) | 123 | 0.523 |

| | **IWR-ON Mean (SD)** | **IWR-OFF Mean (SD)** | **W** | ***p*** |
|---|---|---|---|---|
| Drift (abs(ΔCr (ppm)) | 0.029 (0.004) | 0.040 (0.016) | 73 | 0.089 |

*Note*. Group mean (standard deviation (SD)) and results of a non-parametric paired samples Wilcoxon signed rank test (top: test versus retest) and between-group Wilcoxon signed rank test (bottom: IWR-ON versus IWR-OFF) are presented. Frequency drift was determined based on the creatine signal at 3.02 ppm in each average. Mean drift was calculated for each participant as the average of the absolute value of change in the creatine signal across 4 experiments (top: test versus retest) or across all 8 experiments (bottom: IWR-ON versus IWR-OFF). Drift was assessed before applying frequency correction in Osprey. IWR = interleaved water referencing.

**Table B2.2.** Within-subject CVs for GABA+ with IWR-ON on versus IWR-OFF

| **IWR-ON** | | | | | |
|---|---|---|---|---|---|
| | | **MEGA-14** | **MEGA-20** | **HERMES-14** | **HERMES-20** |
| 0.4 ppm knot | 1to1GABA | **22.9%** (4.8-32.0%) | **13.2%** (9.2-16.2%) | **31.9%** (13.2-43.2%) | **13.3%** (6.9-17.5%) |
| | 1to1GABAsoft | **17.9%** (NaN-26.5%) | **9.2%** (5.9-11.6%) | **27.6%** (NaN-40.1%) | **16.4%** (NaN-25.3%) |
| | 3to2MM | **18.1%** (5.9-25.0%) | **9.0%** (6.3-11.0%) | **10.8%** (4.4-14.7%) | **11.4%** (7.3-14.4%) |
| 0.55 ppm knot | 1to1GABA | **5.7%** (4.2-6.9%) | **7.2%** (4.9-9.0%) | **29.6%** (20.0-36.8%) | **10.7%** (5.3-14.2%) |
| | 1to1GABAsoft | **8.6%** (4.0-11.5%) | **8.9%** (NaN-12.7%) | **15.9%** (11.7-19.1%) | **13.7%** (7.3-17.9%) |
| | 3to2MM | **7.5%** (NaN-11.4%) | **10.2%** (6.7-12.8%) | **8.4%** (4.4-11.0%) | **5.8%** (3.9-7.2%) |
| Gannet | | **19.9%** (3.6-28.0%) | **21.8%** (NaN-31.2%) | **22.1%** (8.7-30.0%) | **30.8%** (15.7-40.6%) |
| **IWR-OFF** | | | | | |
| | | **MEGA-14** | **MEGA-20** | **HERMES-14** | **HERMES-20** |
| 0.4 ppm knot | 1to1GABA | **19.0%** (NaN-28.2%) | **17.2%** (12.3-21.0%) | **36.7%** (25.7-45.0%) | **22.6%** (7.5-31.0%) |
| | 1to1GABAsoft | **15.3%** (NaN-22.6%) | **24.7%** (14.7-31.7%) | **32.1%** (16.1-42.4%) | **18.1%** (13.5-21.8%) |
| | 3to2MM | **15.3%** (9.4-19.5%) | **10.7%** (8.0-12.9%) | **11.8%** (3.4-16.4%) | **11.4%** (7.5-14.2%) |
| 0.55 ppm knot | 1to1GABA | **9.8%** (NaN-14.6%) | **10.7%** (NaN-15.6%) | **28.7%** (19.5-35.7%) | **19.5%** (NaN-28.1%) |
| | 1to1GABAsoft | **9.4%** (4.9-12.4%) | **13.1%** (8.3-16.5%) | **21.3%** (11.9-27.7%) | **11.4%** (NaN-17.7%) |
| | 3to2MM | **10.3%** (3.5-14.1%) | **9.7%** (5.8-12.5%) | **13.3%** (4.9-18.1%) | **6.0%** (3.5-7.8%) |
| Gannet | | **15.5%** (5.6-21.2%) | **28.5%** (14.8-37.5%) | **28.8%** (13.6-38.4%) | **25.7%** (11.1-34.6%) |

*Note*. Within-subject coefficients of variation (CVs) and 90% confidence intervals (CIs) were calculated according to the root mean square method described by (Bland, 2006: https://www-users.york.ac.uk/~mb55/meas/cv.htm). This CI formula requires taking the square root of the calculated lower and upper bounds; in several cases, the formula failed because it required taking the square root of a negative number. The lower CI bound is listed in NaN in these cases. See Bland (2006) for details.

**Table B2.3.** Within-subject CVs for GSH with IWR-ON versus IWR-OFF

| | | | |
|---|---|---|---|
| **IWR-ON** | | | |
| | | **HERMES-14** | **HERMES-20** |
| Osprey | 0.4 ppm knot | **9.2%** (5.8-11.6%) | **11.3%** (7.9-14.0%) |
| | 0.55 ppm knot | **15.5%** (9.4-19.9%) | **13.6%** (8.7-17.2%) |
| Gannet | | **34.3%** (13.6-46.5%) | **37.8%** (NaN-58.1%) |
| **IWR-OFF** | | | |
| | | **HERMES-14** | **HERMES-20** |
| Osprey | 0.4 ppm knot | **19.4%** (12.2-24.6%) | **22.3%** (NaN-31.7%) |
| | 0.55 ppm knot | **16.8%** (10.6-21.2%) | **26.4%** (8.0-36.5%) |
| Gannet | | **38.0%** (NaN-54.0%) | **45.2%** (29.6-56.6%) |

*Note*. Within-subject coefficients of variation (CVs) and 90% confidence intervals (CIs) were calculated according to the root mean square method described by (Bland, 2006: https://www-users.york.ac.uk/~mb55/meas/cv.htm). This CI formula requires taking the square root of the calculated lower and upper bounds; in several cases, the formula failed because it required taking the square root of a negative number. The lower CI bound is listed in NaN in these cases. See Bland (2006) for details.

**Section B3. GABA+ Within-Subject Reproducibility with Gannet**

Gannet Processing

We additionally processed all MRS data using Gannet (v 3.2.1)[4] for comparison with our prior test-retest work. We applied the default Gannet pipeline, including robust spectral registration for frequency and phase correction of the individual transients, zero-filling to 32,000 data points, and 3 Hz exponential line broadening. We then fit GABA+ and GSH models to the data. The Gannet model for GABA-edited spectra applies three Gaussian peaks to model GABA+ and Glx (i.e., the glutamate and glutamine doublet) between 2.79 and 4.1 ppm. The Gannet model for GSH-edited HERMES spectra models data between 2.25 and 3.5 ppm and applies a five-Gaussian model, in which one Gaussian is assigned to model the GSH signal and the remainder models the complex co-edited aspartyl multiplet at ~2.6 ppm. In all cases, Gannet uses a four-parameter curved baseline function containing linear and quadratic terms. Water reference data were quantified using a Gaussian-Lorentzian lineshape model. As with the Osprey processing, we quantified metabolite levels with respect to the unsuppressed water scan, and no further relaxation or tissue segmentation corrections were applied.

Data quality were generally good following Gannet processing (Figure B3.1). To match the Osprey analyses, we excluded the same three datasets as described in the main text (i.e., one MEGA-14 and two HERMES-20). We excluded one additional HERMES-14 dataset due to poor alignment of sub-spectra in Gannet (though this same problem did not occur with Osprey).

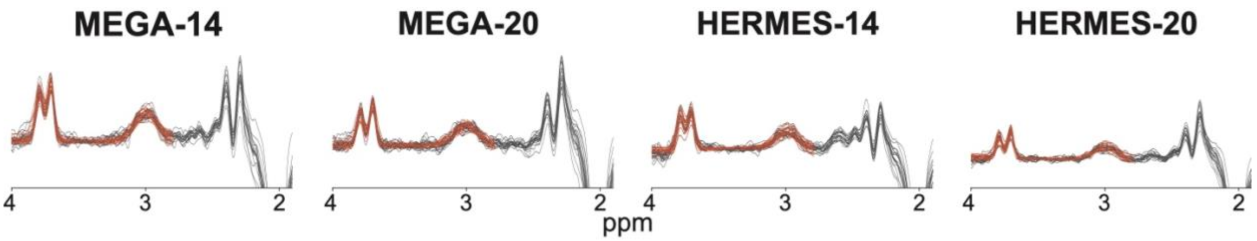


**Fig B3.1. GABA+ Spectra Processed using Gannet.** GABA-edited MEGA-PRESS and HERMES spectra (gray) and model fits (orange) are shown for each participant for the test condition. All spectra are shown from 1.9-4.0 ppm.

Within-Subject Reproducibility of GABA+ Following Gannet Modeling

Within-subject CVs for GABA+ metabolite values resulting from Gannet processing are presented in Table B3.1 and Figure B3.2-B3.3. Within-subject CVs were generally higher (i.e., poorer test-retest reproducibility) for Gannet compared with GABA+ Osprey.

**Table B3.1.** CVs, ICCs, and Pearson r for GABA+ Following Gannet Modeling

| | MEGA-14 | MEGA-20 | HERMES-14 | HERMES-20 |
|---|---|---|---|---|
| Within-subject CV | 17.7% (10.9-22.6%) | 25.4% (16.8-31.7%) | 25.5% (17.4-31.6%) | 28.6% (19.9-35.2%) |
| ICC | 0.07 | 0.02 | -0.27 | -0.14 |
| Pearson r | 0.08 | 0.02 | -0.29 | -0.13 |

*Note*. Within-subject coefficients of variation (CVs) and 90% confidence intervals (CIs) were calculated according to the root mean square method described by (Bland, 2006: https://www-users.york.ac.uk/~mb55/meas/cv.htm). Single-rater, absolute-agreement, two-way mixed effect ICCs[1,2] were calculated using the *irr* package[3]. All ICC values were < 0.50, indicating poor reliability[2]. Pearson correlation coefficients were calculated between test and retest GABA+ estimates. No correlations were statistically significant.

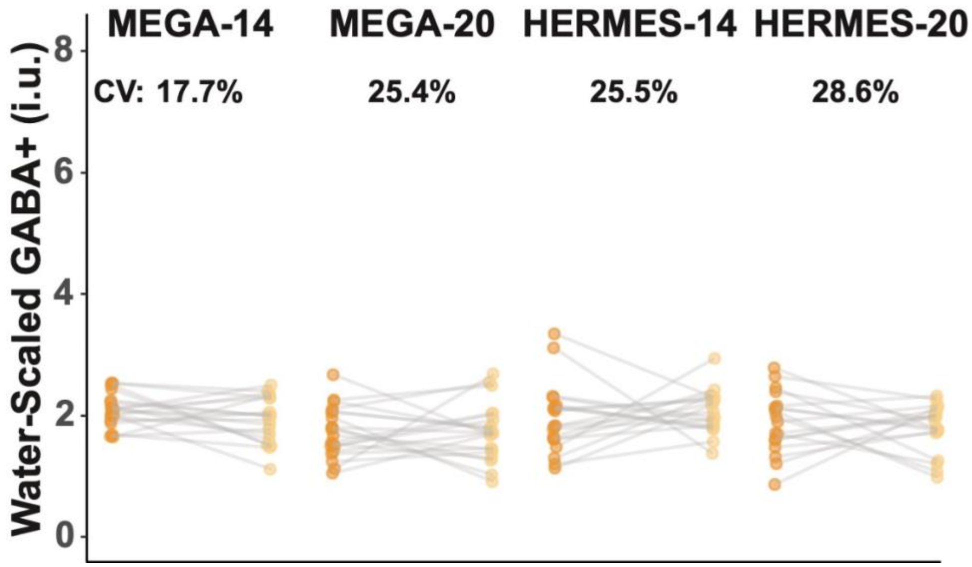


**Fig B3.2. Within-Subject GABA+ values with Gannet.** GABA+ values resulting from Gannet processing are shown for the test (dark orange) and retest (light orange) conditions. Boxplots represent whole-group values, and each point and connecting gray line represents one participant. Within-subject %CVs are listed above each subplot.

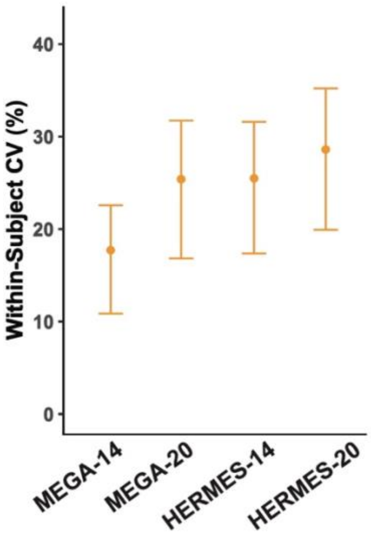


**Fig B3.3. Within-subject CVs for GABA+ with Gannet.** Within-subject %CVs and 90% confidence intervals are shown for each acquisition and post-processing condition.

### <u>Section B4</u>. GSH Within-Subject Reproducibility

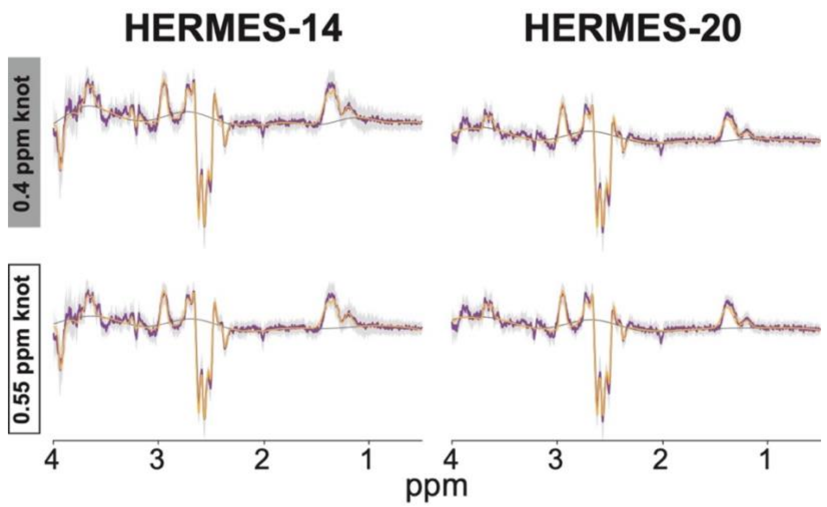


**Fig B4.1. Group Mean GSH Spectra with Osprey.** Average GSH-edited MEGA-PRESS and HERMES spectra (purple), 95% confidence interval (gray shading) and model fits (yellow) are shown for all participants for the test condition. Spectra are shown from 0.5-4 ppm.

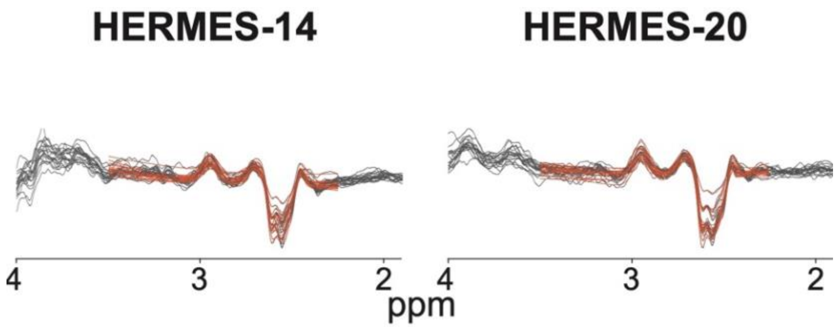


**Fig B4.2. Group Mean GSH Spectra with Gannet.** GSH-edited HERMES difference spectra (gray) and model fits (orange) are shown for each participant for the test condition. Spectra are shown from 1.9-4 ppm.

**Table B4.1.** Within-subject CVs for GSH

| | | HERMES-14 | HERMES-20 |
|---|---|---|---|
| Osprey | 0.4 ppm knot | 15.2% (10.4-18.7%) | 12.9% (10.3-15.0%) |
| | 0.55 ppm knot | 16.2% (12.5-19.2%) | 17.3% (11.6-21.5%) |
| Gannet | | 36.1% (22.5-45.8%) | 41.2% (24.3-53.0%) |

*Note*. Within-subject coefficients of variation (CVs) and 90% confidence intervals (CIs) were calculated according to the root mean square method described by (Bland, 2006: https://www-users.york.ac.uk/~mb55/meas/cv.htm).

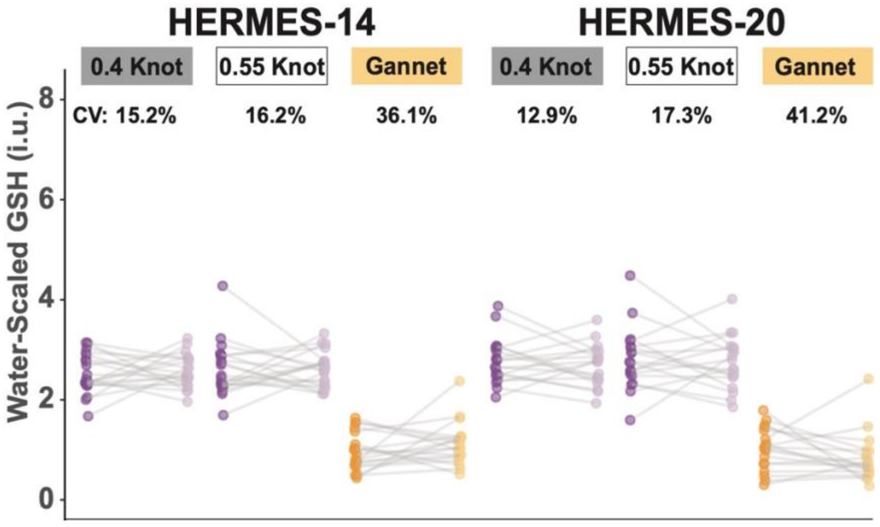


**Fig B4.3. Within-Subject GSH values.** GSH values are shown for the test (dark color) and retest (light color) conditions. Each point and connecting gray line represents one participant. Purple indicates Osprey results and orange indicates Gannet results. Within-subject %CVs are listed above each subplot.

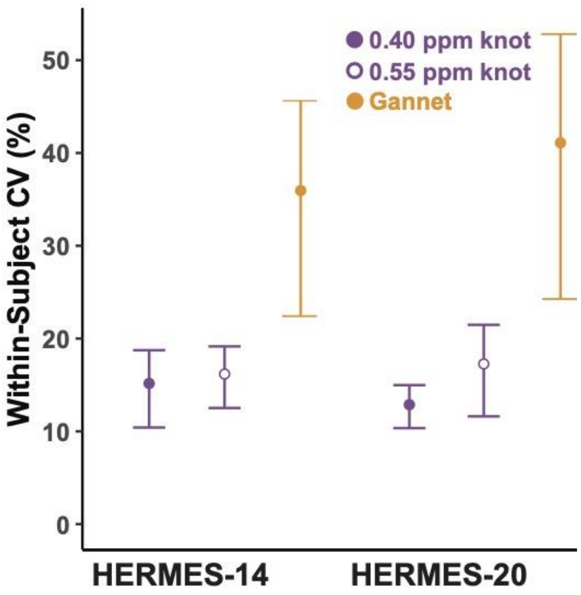


**Fig B4.5. Within-subject CVs for GSH.** Within-subject %CVs and 90% confidence intervals are shown for each acquisition and Osprey (purple) or Gannet (orange) post-processing condition.

**Table B4.2.** ICCs for GSH

| | | HERMES-14 | HERMES-20 |
|---|---|---|---|
| Osprey | 0.4 ppm knot | -0.13 | 0.32 |
| | 0.55 ppm knot | 0.02 | 0.02 |
| Gannet | | 0.01 | 0.18 |

*Note*. Single-rater, absolute-agreement, two-way mixed effect ICCs[1,2] were calculated using the *irr* package[3]. All ICC values were < 0.50, indicating poor reliability[2].

**Table B4.3.** Pearson Correlation Coefficients for GSH

| | | **HERMES-14** | **HERMES-20** |
|---|---|---|---|
| Osprey | 0.4 ppm knot | -0.12 | 0.31 |
| | 0.55 ppm knot | 0.02 | 0.02 |
| Gannet | | 0.01 | 0.19 |

*Note*. Pearson correlation coefficients were calculated between test and retest GSH estimates. No correlations were statistically significant.
